# Development and testing of a novel Killer-Rescue self-limiting gene drive system in *Drosophila melanogaster*

**DOI:** 10.1101/680629

**Authors:** Sophia H. Webster, Michael R. Vella, Maxwell J. Scott

## Abstract

We report the development and laboratory testing of a novel Killer-Rescue (K-R) self-limiting gene drive system in *Drosophila melanogaster.* This K-R system utilizes the well-characterized Gal4/UAS binary expression system and the Gal4 inhibitor, Gal80. Three killer (K) lines were tested; these used either an autoregulated UAS-Gal4 or UAS-Gal4 plus UAS-hid transgene. One universal rescue (R) line was used, UAS-Gal80, to inhibit Gal4 expression. The K lines are lethal and cause death in the absence of R. We show that Gal4 RNA levels are high in the absence of R. Death is possibly due to transcriptional squelching from high levels of Gal4. When R is present, Gal4 activation of Gal80 would lead to inhibition of Gal4 and prevent overexpression. With a single release ratio of 2:1 engineered K-R to wildtype, we find that K drives R through the population while the percent of wild type individuals decreases each generation. The choice of core promoter for a UAS-Gal4 construct strongly influences the K-R system. With the strong *hsp70* core promoter, K was very effective but was quickly lost from the population. With the weaker DSCP core promoter, K persisted for longer allowing the frequency of individuals with at least one copy of R to increase to over 98%. This simple gene drive system could be readily adapted to other species such as mosquito disease vectors for driving anti-viral or anti-parasite genes.

**Significance:** Here we report the development and testing of a novel self-limiting gene drive system, Killer-Rescue, in *Drosophila melanogaster*. This system is composed of an auto-regulated Gal4 Killer (K) and a Gal4-activated Gal80 Rescue (R). Overexpression of Gal4 is lethal but in the presence of R, activation of Gal80 leads to much lower levels of Gal4 and rescue of lethality. We demonstrate that with a single 2:1 engineered to wildtype release, more than 98% of the population carry R after eight generations. We discuss how this Killer-Rescue system may be used for population replacement in a human health pest, *Aedes aegypti*, or for population suppression in an agricultural pest, *Drosophila suzukii*.

## Introduction

Gene drive systems have been proposed as a species-specific genetic method to control insect pests with the end goal of either population replacement or population suppression. These systems have the potential to reduce the impact of human disease vectors or agricultural pests at the individual, community, and global scale. However, some of the recently developed “limitless” systems pose challenges such as introgression into nontarget populations or movement across political borders (1). Instead, several self-limiting gene drive systems have been proposed and/or developed including Daisy-Chain gene drive (1), one or two-locus underdominance (2) and toxin-antidote systems including *Medea* (3), CleaveR (4) and Killer-Rescue (K-R) (5). While some systems, such as Daisy-Chain (1), are theoretical at this time, proof-of-principle experiments with others such as *Medea* (6) and CleaveR (4) have been performed in the model insect *Drosophila melanogaster.* With toxin-antidote systems such as *Medea*, genes of interest linked to the antidote will spread through a population while insects that inherit the toxin alone die. Although successful, the complexity of the *Medea* system has made it difficult to develop for other insects such mosquito disease vectors.

Presented here is a simple and novel toxin-antidote K-R system (5) in *D. melanogaster*. The K-R system utilizes the well-characterized Gal4/UAS binary expression system (7,8) and the Gal4 inhibitor, Gal80. Death is due to overexpression of Gal4 which is rescued by Gal4 activation of the Gal80 inhibitor in flies that have both UAS-Gal4 (K) and UAS-Gal80 (R) transgenes. In one drive experiment over 98% of the flies carry at least one copy of R with a single 2:1 engineered: wildtype release. We show that, as predicted through modeling (5), K drops out of the population over time as it drives the rescue -and linked cargo-each generation. R is also expected to eventually drop out of the population if it has any associated fitness costs and there is migration of wildtype individuals into the release area (5). This simple drive system should be easy to adapt to other insects as the key components, the yeast Gal4 and Gal80 proteins, function in several species (9, 10, 11, 12).

## Results

### Generation of a self-limiting Killer-Rescue gene drive

To create a killer-rescue gene drive in *D. melanogaster*, we designed constructs and made transgenic lines with (1) a Gal80 rescue inserted on the third chromosome and (2) a Gal4 killer inserted on the second chromosome using the phiC31 system for site-directed integration. For the killer (K) we tested an autoregulated UAS-Gal4, either alone or in combination with a Gal4-activated *head involution defective* (*hid*) gene (UAS-hid) (**Fig. 1**). An autoregulated tetracycline transactivator (tTA) transgene has been widely used a conditional lethal system in insects. Death is thought to be due to a general interference in gene expression due to high levels of the tTA transcription factor. Similarly, we reasoned that overexpression of Gal4 would be lethal. Activation of *hid* expression by Gal4 would cause lethality due to induction of apoptosis. As the core promoter can have a significant impact on the level of gene activation by Gal4 (13), we evaluated different core promoters with the Gal4/UAS binary system. All of the systems have the same Gal80 rescue with the strong *D. melanogaster hsp70* core promoter.

**Figure 1.**
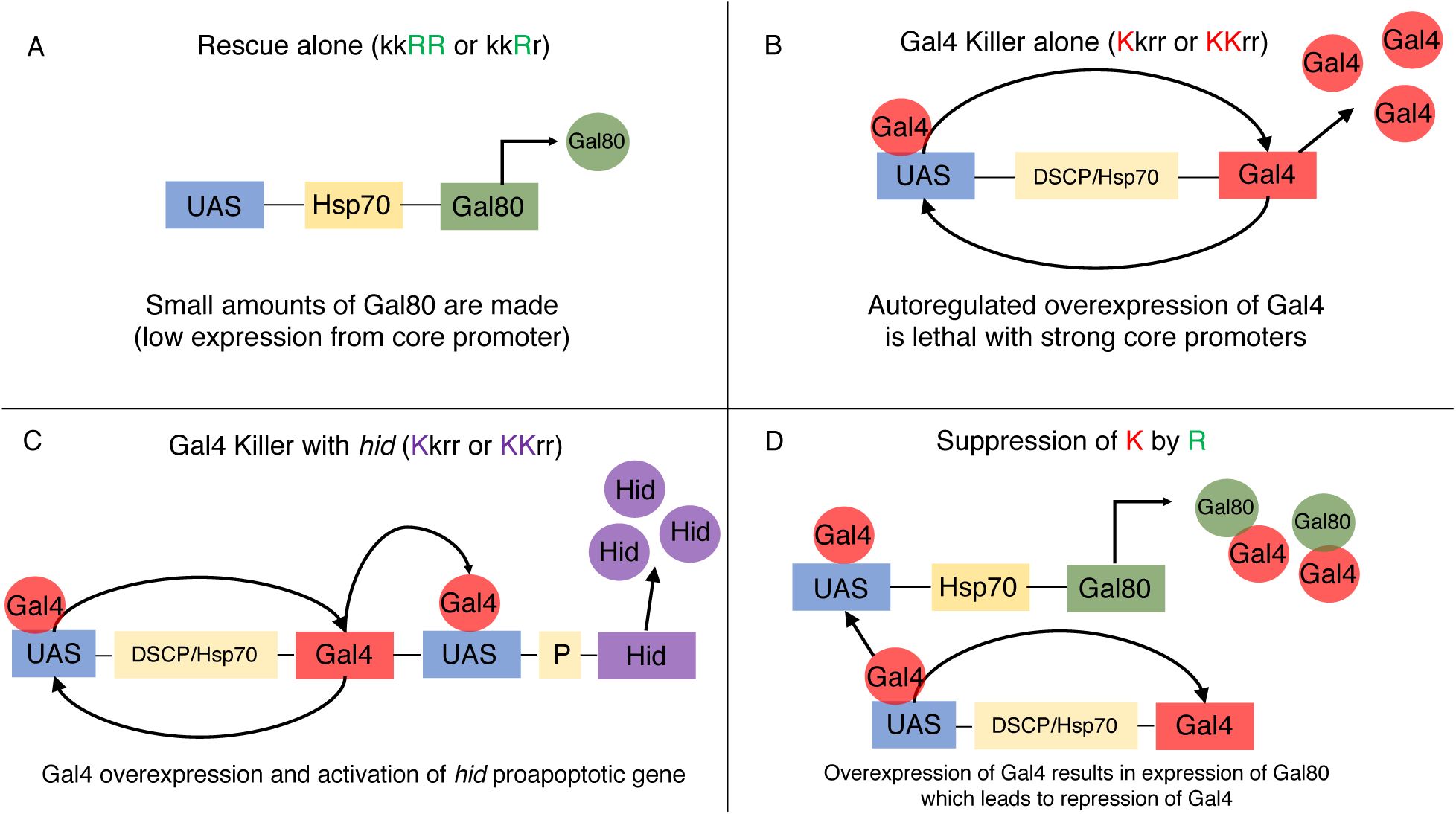
A Gal4-based Killer-Rescue (K-R) self-limiting gene drive system. (A) When the rescue, UAS-Hsp70-Gal80, is present alone in *D. melanogaster* small amounts of Gal80 are expressed from the Hsp70 core promoter. (B) The Gal4 killer, by itself, results in high levels of autoregulated overexpression of Gal4. This autoregulated overexpression is lethal with strong core promoters such as DSCP and Hsp70. (C) The Gal4 killer is modified by the addition of Hid cell death, proapoptotic, gene with Gal4. (D) When K and R are present together in the fly the system “chases” itself. Overexpression of Gal4 results in expression of Gal80 which in turn leads to repression of Gal4. The lethal phenotype is rescued by Gal80 repression of Gal4 and the flies survive.

By using different core promoters, we were able to modulate the expression of Gal4 and optimize the lethal system. The strong core promoter from the *D. melanogaster hsp70* gene (*DmHsp70*) has been widely used since the development of the Gal4/UAS binary system in Drosophila (7). The Drosophila synthetic core promoter (DSCP) contains optimized versions of the motifs (TATA, Inr, MTE, and DPE) that are thought be important for function (14). Indeed, the DSCP core promoter has been found to promote robust expression with a broad range of enhancers that bind different activator proteins. However, in the specific case of Gal4-driven UAS expression, the *DmHsp70* core promoter yields approximately twofold higher expression levels than the same construct when built with DSCP (14). Therefore, we expected that in the lines carrying a UAS-Gal4 transgene, the level of Gal4 gene expression would be higher when driven with the *DmHsp70* core promoter than with the DSCP. The *hsp70* core promoter was used to ensure that Gal4 activated high levels of expression of Gal80.

The synthetic translational enhancer *syn21* (15) was used in both killer and rescue constructs to increase translation efficiency and thus increase protein production of Gal4 and Gal80. This translational enhancer is a synthetic AT-rich 21-bp sequence made by combining the Cavener consensus sequence (Cavener, 1987) with elements from the *Malacosoma neustria* nucleopolyhedrovirus (MnNPV) polyhedron gene (16).

### K-R system exhibits lethality (K alone) and repression of lethal effects (R present)

To establish K-R strains, a line homozygous for R was used as the recipient for germline transformation with R constructs. We then tested the ability of the rescue UAS-Gal80 line to repress activation of a RedStinger reporter gene crossed with Gal4-drivers. (**Fig. S1, S2**) Repression of red fluorescence was quite clear in flies that carried UAS-Gal80, a Gal4-driver, and UAS-RedStinger. With the DSCP promoter, strains homozygous for K and R were established by selecting for strong fluorescence of the marker genes (**Fig. S3**). Further, one copy of R was able to repress two copies of K (KKRr flies survive). However, we could not establish double homozygous lines, KKRR, with any K that used the *hsp70* promoter. To determine if K was lethal, flies heterozygous for K and R were crossed with wild type. We observed that insects with only K did not survive to adulthood. With the strong *hsp70* promoter, death occurred at the early larval stages but with the DSCP promoter death occurred at the late pupal stage. These results are consistent with the expectation that K constructs built with the stronger *hsp70* core promoter would be particularly effective.

### 2:1 Single Release Population Cage Experiments

Since K alone is lethal and R is able to rescue lethality from K, we performed multigenerational population cage experiments to determine if K could drive R through a population as predicted from models (5). The input for each experiment was 100 K-R (or R alone) flies and 50 wild type flies per bottle (2:1 release ratio). Five biological replicates were set for each release experiment. The input fly line DSCP-Gal4; Gal80 was homozygous for both transgenes (KKRR, where a capital K or R indicates the transgenic allele and a lowercase k or r indicates the wildtype allele). However, the hsp70-Gal4; Gal80 and hsp70-Gal4-hid; Gal80 lines were homozygous for R but heterozygous for K (KkRR) as flies homozygous for K did not survive. The input Gal80 fly line was homozygous (kkRR). These population cage experiments were followed for six generations (Hsp70-Gal4, Hsp70-Gal4-Hid killers and Hsp70-Gal80 rescue alone) or for nine generations (DSCP-Gal4 killer) (with the starting release defined as generation one). The genotypes were calculated each generation by screening fluorescence intensity of adult individuals. Each subsequent generation bottle was set with 150 flies, with genotypes in the same proportion as counted in the total. For the DSCP-Gal4; Gal80 experiment, there are nine possible genotypes in the population after release (KKRR, kkRR, kkRr, KkRr, KKRr, KkRR, Kkrr, KKrr, kkrr (wt)). Of these, Kkrr, KKrr should die and the other seven should survive. For the Hsp70-Gal4; Gal80 and Hsp70-Gal4-Hid; Gal80 experiments there are only five viable genotypes (kkRR, kkRr, KkRr, KkRR, kkrr (wt)) because flies with two copies of K, with or without R, do not survive.

The most successful gene drive cage experiment was with the DSCP-Gal4 killer: within 4 generations over 98% of flies carried at least one copy of R (**Fig. 2A**). In the gene drive experiment with hsp70-Gal4 and hsp70-Gal4-hid the killers are rapidly lost from the population (**Fig. 2C**), which suggests incomplete rescue by R. Nevertheless, all killers were successful in driving R as more than 94% of the flies carried at least one copy of R by the end of the experiment (**Fig. 2A**) and all show loss of wildtype (**Fig. 2C**).

**Figure 2.**
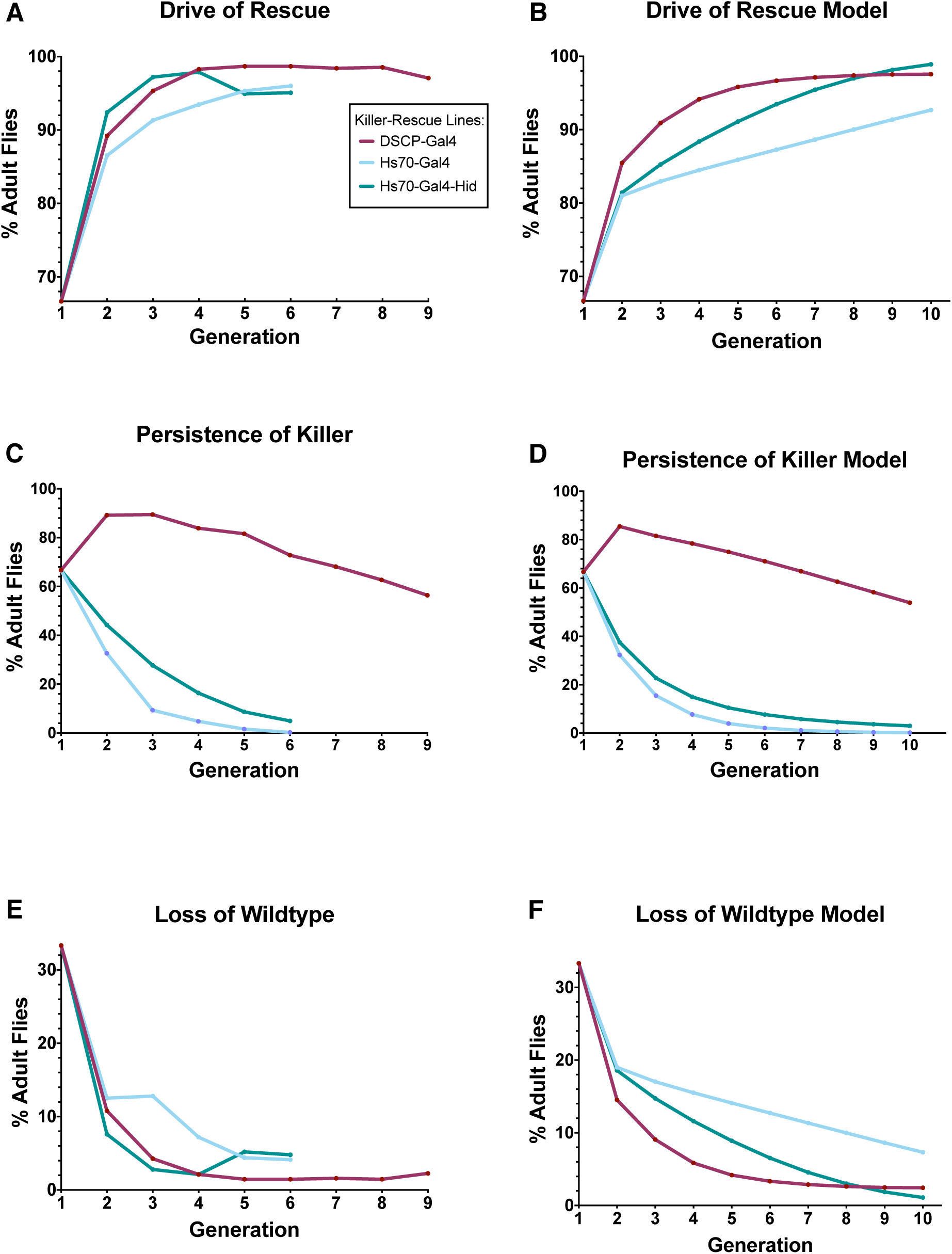
A comparison of K-R gene drive experiments. Starting release (generation 1) of equal numbers males and females, 2:1 engineered: wildtype adult flies (100: 50). Comparing Killer-Rescue gene drive experiments with (A) Drive of Rescue, (B) Persistence of Killer, and (C) Loss of Wildtype. In all experiments, the killer drove the rescue to more than 94% of flies. All three Killer-Rescue fly lines similarly repress the wildtype, and the DSCP-Gal4 K-R is able to maintain suppression of the wildtype, while the killer is maintained in the population at a high frequency through generation nine. In K-R lines with the hsp70 core promoter, the wildtype population will likely increase after generation six because the killer is no longer present to suppress the wild population and drive the rescue into the population.

For the DSCP-Gal4 killer, because one copy of R was able to repress one and two copies of the killer, the killer transgenes are largely maintained in this drive and still present in 56% of the flies by generation nine (**Fig. 2C**). There was a steady decrease in the proportion of flies that are homozygous for K but heterozygous for R (KKRr) with each generation (**Fig. 3A**). Further, with each generation there was a steady increase in the proportion of the population that lack K but are homozygous for R (kkRR). In the experiments with the hsp70-Gal4 and hsp70-Gal4-hid killers, there was a more rapid increase in the proportion that were kkRR (**Fig. 3 C,E**). The proportion of the population that were KkRr decreased rapidly after the first generation, in contrast to what was observed with the DSCP-Gal4 killer. The UAS-Gal80 transgene (R) appears to have only a small fitness cost as the proportion of flies homozygous for R shows a small decrease each generation that is matched by an increase in wildtype (**Fig. 3G**).

**Figure 3.**
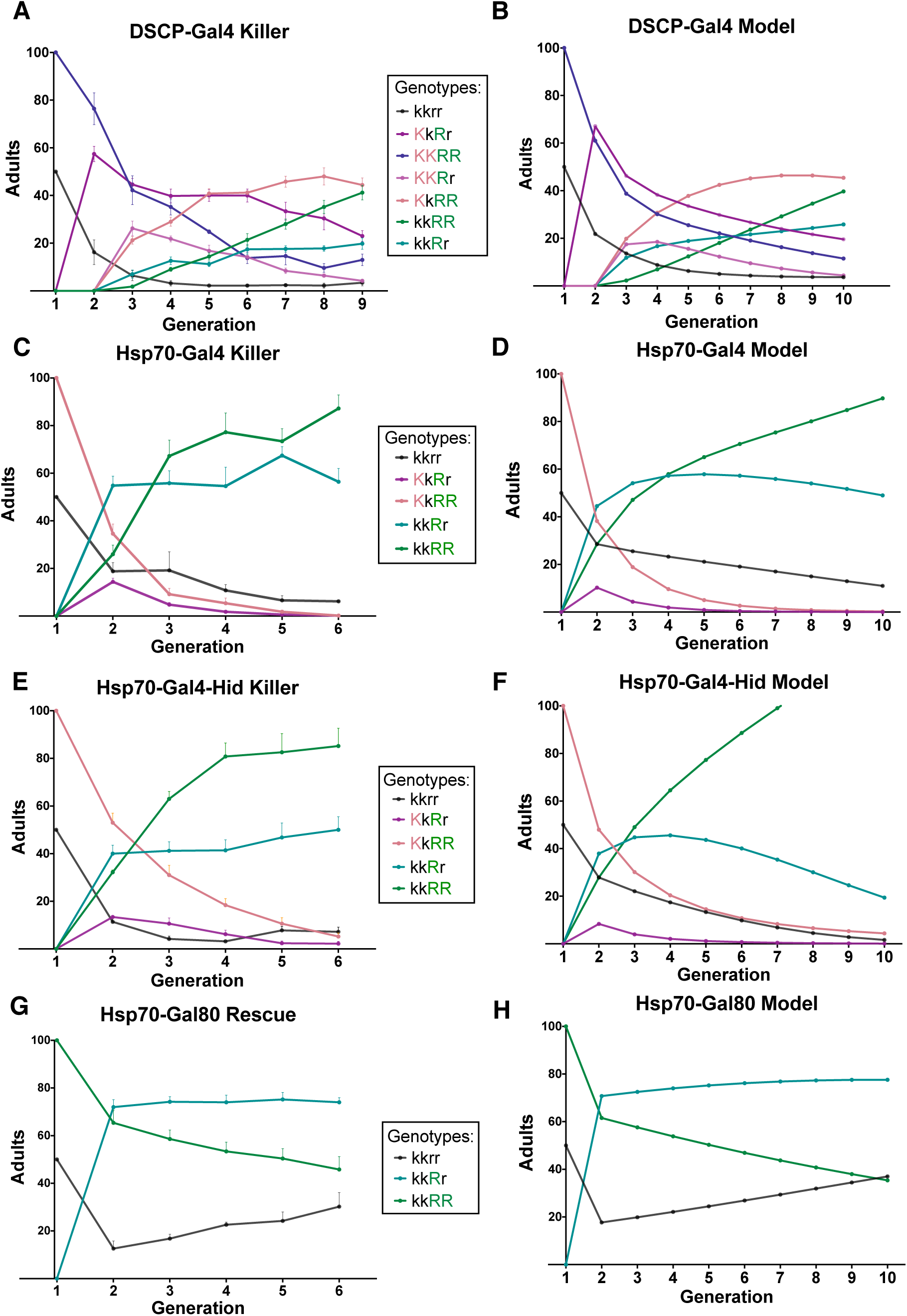
Gene Drive Release Experiments and Model Simulations. Starting release (generation 1) of equal numbers males and females, 2:1 engineered: wildtype adult flies (100: 50). In (A), (C), (E), and (G), the plots display an average of the genotypes, from 5 replicates, present in each generation of the gene drive. The Hsp70-Gal80 rescue (G) was tested alone to determine fitness costs associated with the rescue by itself. The three different killers were tested in (A) DSCP-Gal4, (C) hsp70-Gal4 and (E) hsp70-Gal4-hid Killer. Modeling (B,D,F,H) fitness parameters for each genotype were calculated by minimizing the error sum of squares. Equal numbers of males and females were added back to the cage each generation after counting the genotypes to maintain the population at 150 adults every generation. Error bars represent the SEM (+/−) of the five averaged biological replicates performed in each gene drive experiment.

### Evaluating K-R Dynamics Using Mathematical Modeling

A mathematical model was used to calculate expected population dynamics over time and can be used to estimate fitness parameters. The fitness parameters for genotypes that do not survive to adulthood were set to 0 (eg. Kkrr and KKrr). The remaining fitness parameters depend on the strength of the promoter (Hsp70 or DSCP), and presence or absence of hid, and are thus different for each K-R gene drive experiment. The fitness parameters that minimized the error sum of squares were used for model simulations (**Fig. 2, 3; Table S1**). The fitness values from the Hsp70-Gal80 rescue alone suggest that R induces a minor fitness cost. The highest consistency between model and data indicate somewhat lower fitness than wildtype when homozygous (0.87), but not when heterozygous (1.0), though a range of low fitness costs are also consistent with data (**Fig. S4).** The fitness estimates from the other experiments show that R is most effective at rescuing DSCP-Gal4 (KkRr = 0.77 (relative fitness) and less effective at rescuing Hsp70-Gal4 (KkRr = 0.18) and Hsp70-Gal4-hid (KkRr = 0.15). For these experiments, the model trajectories are not as consistent with the empirical data, and the fitness estimates do not completely follow the expectation that each killer (K) allele induces a fitness cost which can be partially mitigated by each copy of rescue (R). This could be because the model does not account for stochasticity or complex interactions between genotypes that could lead to changes in fitness over time.

### Gal4 Expression and Gal80 Repression of Gal4

We next performed qRT-PCR experiments to quantify the levels of Gal4 and Gal80 expression during late embryo (18-20 hour) and mid-pupal (48 hour) stages of development. We chose these two life stages because they appear to be critical time points in the fly where we observed lethality from a killer. In the lines with the strong *hsp70* core promoter, we observed that first instar larvae containing only the killer (Kkrr) died. With the DSCP-Gal4 killer line, flies with one copy of the killer, Kkrr, die during late pupal development. In DSCP-Gal4 embryos, GAL4 expression is 35-fold lower in the presence of R than in the absence (**Fig. 4A**). In pupae, in the absence of R, GAL4 RNA levels are high, nearly three times higher than in embryos. Repression by GAL80 is significant, as GAL4 levels are much lower in the presence of R (**Fig. 4B**). Although GAL4 RNA levels are higher than observed in embryos this does not appear to be deleterious as flies with two copies of K and one or two copies of R (KKRR and KKRr) survive and are fertile. Hsp70-Gal4 embryos with only K express nearly 10-fold greater Gal4 than embryos with K and two copies of R (**Fig. 4C**). Hsp70-Gal4-hid embryos with only K express Gal4 nearly 90-fold greater than embryos that carry K and two copies of R (KkRR) (**Fig. 4D**).

**Figure 4.**
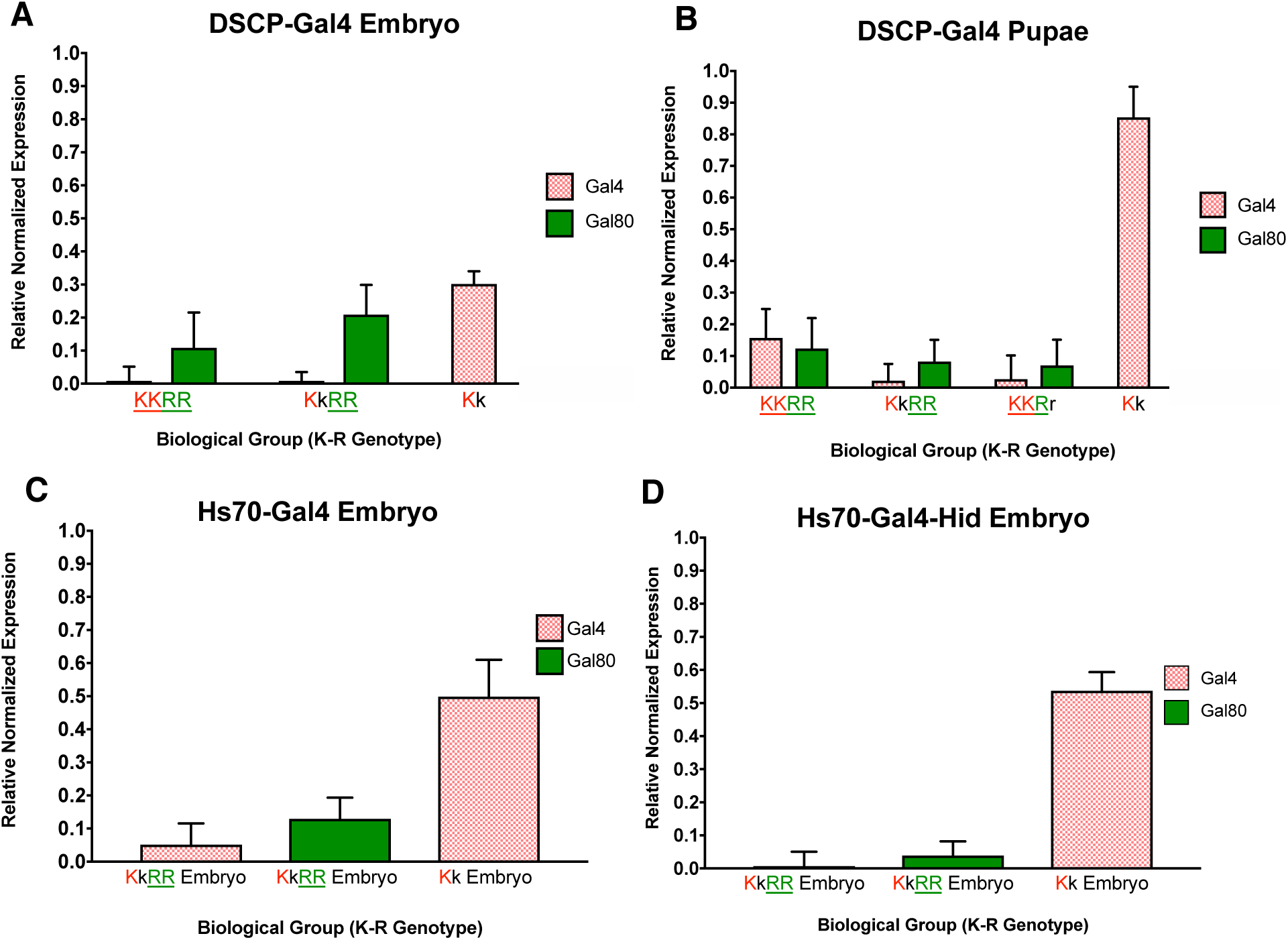
Gal80 (R) and Gal4 (K) expression in embryos and pupae. The relative normalized expression determined by qRT-PCR of Gal4 and Gal80 in 18-20 hour old embryos and 48 hour old pupae from each line are shown. For each sample four biological replicates were collected and each of these biological replicates are averaged from four technical replicates.

## Discussion

Gene drive systems have great potential for insect pest population replacement or suppression (17). We have described a novel Killer-Rescue (K-R) gene drive system in *D. melanogaster* that successfully drove R, the rescue transgene, in multigenerational population cage experiments. This particular K-R system, constructed with Gal4 and Gal80 transgenes, should be readily transferable to a number of insect pests such as mosquito disease vectors. An advantage of this system is that Gal4 expression levels can be easily modulated to achieve the desired properties for drive. The number of Gal4 binding sites (UAS), the core promoter, presence of a 5’UTR translation enhancer, a 3’UTR RNA transport signal and the type of transcription terminator have all been shown to influence overall expression with the Gal4/UAS binary system (13).

High expression levels of the transcription transactivator (tTA) are lethal in a number of insect species, including *D. melanogaster* (18), the New World screwworm *Cochliomyia hominivorax* (19), the mosquito *Aedes aegypti* (20) and the silk moth *Bombyx mori* (21). Similarly, the K-R system developed in this study uses an autoregulated UAS-Gal4 transgene as the killer, which we showed leads to high levels of Gal4 RNA. Death is possibly due to a general interference in transcription, as hypothesized for tTA overexpression (20). Gal80 represses the activity of Gal4 by binding to the transcription activation domain and preventing the recruitment of transcriptional machinery by Gal4 (11,12). By creating an autoregulated UAS-Gal4 (killer) and Gal4-activated UAS-Gal80 (rescue) system, the presence of both killer and rescue in the fly will prevent Gal4 over-production and thus suppress lethality. Indeed, we showed that Gal4 expression is much higher in embryos or pupae with only K compared to those with K and R. Additionally, to our knowledge, this is the first demonstration that the widely used Gal4/UAS system can be lethal to *D. melanogaster*.

The multi-generational gene drive experiments demonstrated the importance of choice of core promoter with the GAL4 overexpression system. Similarly, we had previously shown that the core promoter is important for the tTA overexpression system (22). To minimize differences in gene expression due to location, all killer transgenes were inserted at the same position in the genome. In the absence of R, Gal4 RNA levels were approximately two-fold higher in embryos with the *hsp70* promoter than the *DSCP* promoter driving expression. This is consistent with previous observations that the *hsp70* promoter is about twice as active as the DSCP promoter with the Gal4/UAS system (14). The high levels of Gal4 expression could explain why rescue was incomplete with killer constructs that used the strong *hsp70* core promoter. In the drive experiments, we found that there was a rapid decrease in the proportion of flies carrying a *hsp70-Gal4* or *hsp70-Gal4-hid* transgene such that flies with killer were present in less than 5% of the population after 5 generations. In contrast, the DSCP-Gal4 killer is maintained in the population at a much higher frequency throughout the drive experiment and at generation 9 is still present in over 56% of the population. Further, because the only flies that survive have at least one copy of the rescue, all the individuals that have R or K-R are contributing to the spread of the rescue. Maintaining some killer alleles in the population over time could lead to more resiliency for maintaining the rescue in the population. For example, after a release, if wildtype insects migrate into the area where a release occurred, the presence of the killer will continue to suppress the wildtype. Eventually, however, the killer would fall out of the population, followed by the rescue, which would no longer have a selective advantage over the wildtype r allele. K-R is temporally self-limiting in this fashion, and the length of time that the population remains transformed could be modulated by the release ratio.

When designing the K-R systems, we expected that the K construct with the *hid* proapoptotic cell death gene would be more lethal than the K constructs without it. However, the gene drive experiments with *hsp70*-Gal4 and *hsp70*-Gal4-*hid* killers showed similar kinetics. The *hsp70*-Gal4-*hid* killer showed slightly higher levels of persistence in gene drive experiments (5% versus 0.3% after generation five) and more rapidly drove the rescue above 95%. The higher persistence could indicate Gal4 levels are lower and thus more efficiently rescued by *hsp70-Gal80*. In the *hsp70*-Gal4-*hid* transgene there are two UAS binding sites to which Gal4 will bind, thus dividing the binding of Gal4 between the UAS site which will produce more Gal4 and to the UAS site which will induce *hid* expression. This could result in decreased Gal4 expression, which in turn would reduce *Gal80* levels. We did observe lower Gal80 RNA levels in *hsp70*-Gal4-*hid* embryos than *hsp70*-Gal4 embryos (**Fig 4. C, D**). However, Gal4 RNA levels were similar. Nevertheless, the inclusion of a Gal4-activated cell death gene such as *hid* could be beneficial if resistance develops to Gal4 overexpression.

The Killer-Rescue (K-R) gene drive system was originally proposed as a means for driving anti-viral/anti-pathogen genes through a wild population of mosquito disease vectors (5). The modeling demonstrated that with a fitness cost of only 10% for each transgene, the proportion of mosquitoes expected to transmit a virus (i.e. lack rescue and linked anti-pathogen gene), decreased below 5% by 40 generations after a single 2:1 release of engineered: wildtype mosquitoes. Modeling of our system suggests that the rescue alone (kkRr and kkRR) has less than a 10% fitness cost for each transgene. It should be relatively straightforward to adapt the Gal4-Gal80 K-R system for *Ae. aegypti* as germline transformation is routine (23,24,25) and the *D. melanogaster hsp70* core promoter is functional in this species (26). Further, the Gal4/UAS binary system has been used to control gene expression in *Ae. aegypti* (10). *Ae. aegypti* is a vector for multiple viruses, including Zika, dengue, and chikungunya; and causes global morbidity and mortality to millions of people each year (27,28). Recently, engineered resistance to Zika virus in *Ae. aegypti* has been achieved through expression of a polycistronic cluster of synthetic small RNAs that are designed to target the Zika virus genome (29). A cargo such as the one described for Zika resistance (29) could be linked to the rescue in Killer-Rescue and transgenic insects released to transform a wild population of mosquitoes. Similar genetic engineering strategies could be used to engineer mosquitoes that are resistant to other arboviruses such as dengue and chikungunya.

An agricultural pest such as *Drosophila suzukii* would be an attractive target for K-R as gene constructs developed in the close relative *D. melanogaster* would be expected to function in *D. suzukii*. However, adapting Killer-Rescue for suppression of *D. suzukii* populations is less straightforward as drive will not occur if the rescue plus cargo has a significant negative fitness cost (5). Drive could occur if the cargo has minimal impact on the population at the time of release. Options for this type of cargo are (i) increased susceptibility to a certain chemical that would be applied at a later date, (ii) increased susceptibility to parasitism (novel parasitoids of *D. suzukii* have been described (30)), (iii) decreased overwintering survival by disrupting genes associated with the winter morph (31) and (iv) decreased survival during the warmer days of summer (with an early spring release).

The Gal4-Gal80 K-R system provides an alternative to Cas9-based drive systems such as Cleave and Rescue (4) and homing drives (32). The appeal of the K-R system is its simplicity and self-limiting design, which will limit the spread of transgenes in space and time. In contrast, Cas9-based homing drives have a very low threshold for drive and are not self-limiting. In addition, the rapid development of resistance to Cas9 cleavage can lead to drive breakdown within a few generations (33,34,35,36), and complicated design, such as targeting haploinsufficient genes, is required to overcome this weakness (37). While resistance to Gal4 overexpression could also develop, the K-R system could be easily modified if needed to increase lethality.

## Materials and Methods

### Assembly of K and R Constructs

To generate the rescue (R) plasmid UAS-hsp70-Gal80-PolyUb-EGFP-attB, plasmids pUASTattB (GenBank EF362409.1) and MS-1419 (gift from Marc Schetelig) were used to construct the marker Poly-Ub-EGFP with an attB site. The fragment UAS-hsp70-Gal80-p10 was digested out of pBC SK+ with the enzymes *MluI* and *EcoRV* and ligated into the PolyUb-EGFP vector.

To generate the killer constructs with the hsp70 core promoter, first the plasmid PolyUb-DsRed-attB was constructed from source plasmids pUASTattB (GenBank EF362409.1) and MS-1425 (gift from Marc Schetelig). The insert UAS-hsp70-Gal4-p10 was prepared with source plasmids #1299 and pCasperGal4: pC3G4 DGRC #1224. To add *hid* to the UAS-hsp70-Gal4 construct, the plasmid pBC-eryO-tetO7-P-CG5123-HidCDS (created by Fang Li), was used as a template for the following *hid* PCR primers: For 5’-AAAAA<u>GCGGCCGC</u>GATCCCCGACACCAGACCAACT-3’ and Rev 5’- CCCCC<u>GGATCC</u>GAAGAGAACTCTGAATAGGGAATTGGGA-3’. To generate the less lethal killer construct with DSCP, a nucleotide fragment containing DSCP and the appropriate restriction sites for cloning was synthesized and cloned into the original UAS-hsp70-Gal4 killer construct described above. The synthetic translational enhancer *syn21* (15) was used in killer and rescue constructs to increase protein production of Gal4 and Gal80. Ten copies of the UAS (10x UAS) were used in each construct, as this number of copies has been shown to be optimal for greatest expression without leaky basal expression (13). All killer and rescue constructs were designed with the constitutive *Polyubiquitin* (*PUb*) promoter for strong full-body marker gene expression throughout all life stages. Additionally, all constructs contain an attB site for site-specific integration into an attP site in the Drosophila stock line of choice.

### Fly Rearing and Strains

#### Injections

Plasmid DNA was prepared using Invitrogen Purelink HiPure Midiprep. Injection mixes were prepared to a final concentration of 750 ng/μL and then put through a 0.45μM Millipore Ultrafree-MC Centrifugal Filter (Cat #: UFC30HV00) to remove any additional debris before embryo injections. Over 200 embryos were injected for each construct to establish K and R lines. To establish K lines and increase survival of injected embryos with K, a construct containing Gal80 was included in the injection mix for transient expression of Gal80. The surviving K injected embryos were then mated to the homozygous UAS-Gal80 R fly line so that all G_1_ offspring would have one copy of R to repress lethal effects of K.

#### Crosses to establish transgenic lines

To establish transgenic lines, single pair crosses were set between the injected G_0_ flies and the relevant stock line (**Table S2.**). In all subsequent generations, only transgenic flies were collected and set as mates to make the line homozygous for the transgenes. Flies homozygous for the transgenes were collected based on fluorescence intensity and were made double homozygous for K and R in this manner.

### Gene Drive Experiments

The experiment was designed to simulate a 2:1 single release of engineered to wild type flies, respectively. For the wild type line, attP40 (BDSC Stock # 25709) was used because the attP40 background is the same as that of the UAS-Gal80 rescue. The initial cage was comprised of 50 genetically engineered virgin females, 50 genetically engineered males, 25 attP40 virgin females, and 25 attP40 males. Five biological replicates of each Killer-Rescue gene drive experiment were conducted. 8oz round bottom bottles (Cat # 32-129F) from Genesee Scientific were filled with 75 mL of fly food and after three days the cross was transferred to a second bottle. This second bottle was used to count genotypes for the next generation of offspring. Three days later the flies in the second bottle were removed and discarded. Fifteen days after the second bottle was set, all the adult flies were genotyped by marker screening under the fluorescent scope and sorted into their respective genotypic groupings based on fluorescence intensity (**Fig. S3**). The genotypes were counted and then the correct proportion of each genotype was added to the next generation of flies so that the starting population for each generation did not exceed 150 flies (the initial starting population). The experiment was carried out for six (UAS-Gal80, hsp70-Gal4, hsp70-Gal4-hid) or nine (DSCP-Gal4) generations.

### qRT-PCR

#### Sample Collection

Four replicates of each sample were collected from different parents. For embryos, around 100-150 were collected for each replicate. For pupae, 6 or more were collected for each replicate. On average the total amount (μg) of RNA collected from 25-30 pupae was 70-90 μg, which equates to ∼2.8-3 μg/pupae. On average the total amount (μg) of RNA collected from 100 embryos was 6 μg. 3.5 μg of RNA were used in each reverse transcriptase (RT) reaction.

#### RNA Preparation

Pre-filled tubes containing zirconium beads were used for sample collection (Benchmark Scientific Triple-Pure High Impact Zirconium Beads 1.5 mm Cat # D1032-15). 500 μL of Trizol ® was pipetted into each pre-filled tube and then the samples were directly collected into the Trizol ®. After the tissues were disrupted using the homogenizer (OPS Diagnostic HT mini homogenizer 230V Cat # BM-D1030) for 3-4 minutes at max speed (4000 rpm), they were processed immediately using a phenol-chloroform extraction protocol and Qiagen RNEasy Mini Kit (Cat# 74104). Thermo Fisher Scientific dsDNAse (Cat # EN0771) was added to the purified RNA to remove any contaminating DNA. cDNA was synthesized using the Invitrogen (Cat#18080-400 Invitrogen) SuperScript III First-Strand Synthesis SuperMix kit.

To measure relative transcript levels, qRT-PCR was performed with the cDNA template diluted 1:4 with nuclease-free water then pipetted into quadruplicate wells of a 384 well optical plate (Cat#4309849 Applied Biosystems). Thermo Maxima SYBR Green/Rox qPCR Master Mix 2X (Cat#K0221) was added to 10μM primers to create a master mix, which was then dispensed into wells using a multichannel pipette. The primer sequences for Gal4, Gal80, Rpl32, and 18S rRNA are listed in supplementary (**Table S3**). The qPCR run was performed on a BioRad CFX384 C1000 Touch Thermocycler.

The relative normalized expression, ΔΔCq, was calculated to quantify the expression of Gal4 and Gal80 present in the samples collected. The calculations for the qPCR were performed as follows: (1) the mean Cq for each biological replicate was calculated by averaging the four technical replicates, (2) the standard deviation and standard error of the mean were calculated, (3) the ΔCq was calculated for each reference gene (Rpl32 and 18S rRNA) by taking the difference between the mean Cq for each sample biological replicate and the corresponding mean Cq of the reference gene, and (4) the ΔΔCq for each sample was calculated as 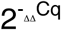.

### Mathematical Modeling

We used a mathematical model to calculate expected population genetics of the cage experiments. The model is discrete-time with non-overlapping generations and assumes random mating and equal sex ratios. The population is normalized every generation, so we track frequencies of each genotype, denoted by subscript *i*. We assume fitness costs are assessed prior to mating. A genotype’s fitness, *w*_*i*_, gives the proportion of individuals of that genotype that enter the mating pool, i.e., survive to adulthood after any viability and embryonic fitness costs are assessed, relative to wildtype fitness of 1. At generation t, the relative proportion of individuals of each genotype entering the next generation, B_i_(t+1), is a function of fitness, current adult frequencies A_i_, and the probability, *P*(*i* |*m,n*), that a mating between female of genotype m and male of genotype n produces offspring of genotype i:

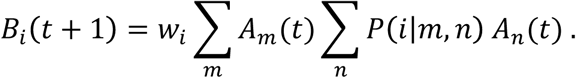

Then, the genotype frequencies of adults in the next generation’s mating pool is:

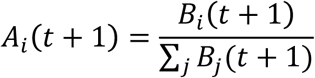

Fitness parameter estimates from the cage experiments were obtained by minimizing the error sum of squares between model and mean experimental genotype frequencies. The fitness values of kkRR and kkRr were estimated using the data from the experimental replicates of Hsp70-Gal80 rescue alone. The remaining four non-zero fitness parameters were estimated for the DSCP-Gal4 replicates. All four non-zero fitness parameters, including those of kkRR and kkRr, were estimated separately for hsp70-Gal4 and hsp70-Gal4-hid killers.

## Supplementary

**Fig S1.**
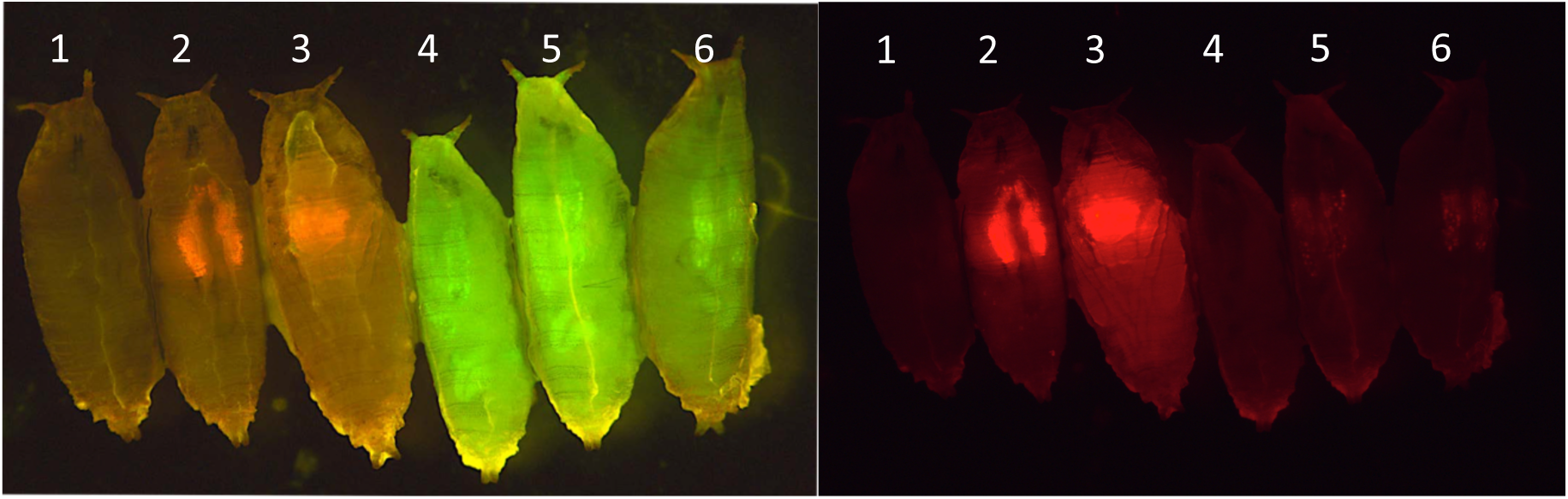
Gal80 repression of the activation of a Redstinger reporter gene by an actin5C-Gal4 driver. Images were taken with the GFP2 filter (left panel) or DsRed filter (right panel) to show green and red fluorescence, respectively. Left to Right: (#1) Wild type, (#2-3) Actin-Gal4 and UAS-RedStinger, (#4) UAS-Gal80 and (#5-6) UAS-Gal80, Actin-Gal4 and UAS-RedStinger. In pupae #5 and #6 the repression of red fluorescence is clearly visible (compare #2,3 with #5,6). The small amount of red fluorescence seen in pupae #5 and #6 was only visible during the pupal stage and not at any other life stage.

**Fig S2.**
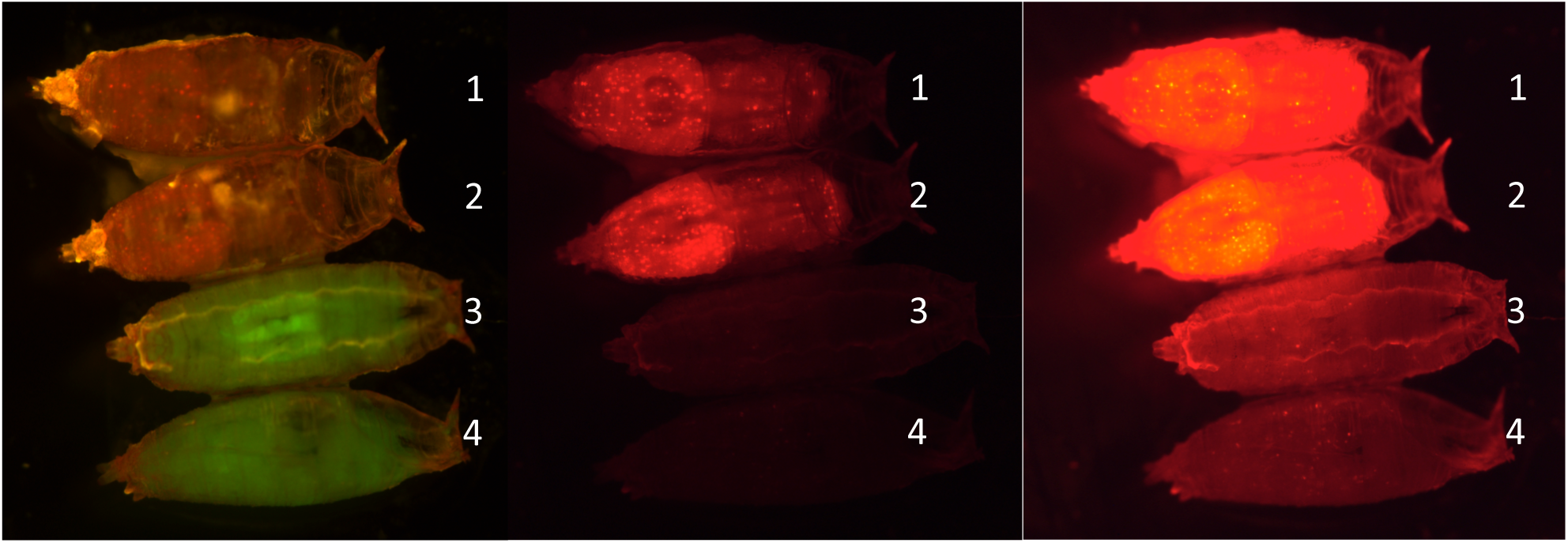
Gal80 repression of the activation of a Redstinger reporter gene by an Elav-Gal4 driver. Images were taken with the GFP2 filter (left panel) or DsRed filter (middle and right panels). Top to Bottom: (#1-2) Elav-Gal4 and UAS-RedStinger, (#3-4) UAS-Gal80 and Elav-Gal4 and UAS-RedStinger. In pupae #3-4 the repression of red fluorescence is clearly visible. There is a small amount of red fluorescence visible as speckles in right panel (#3-4), however DsRed activation is almost entirely repressed by Gal80. The small amount of red fluorescence seen in #3,4 was only visible during the pupal stage and not at any other life stage.

**Fig S3.**
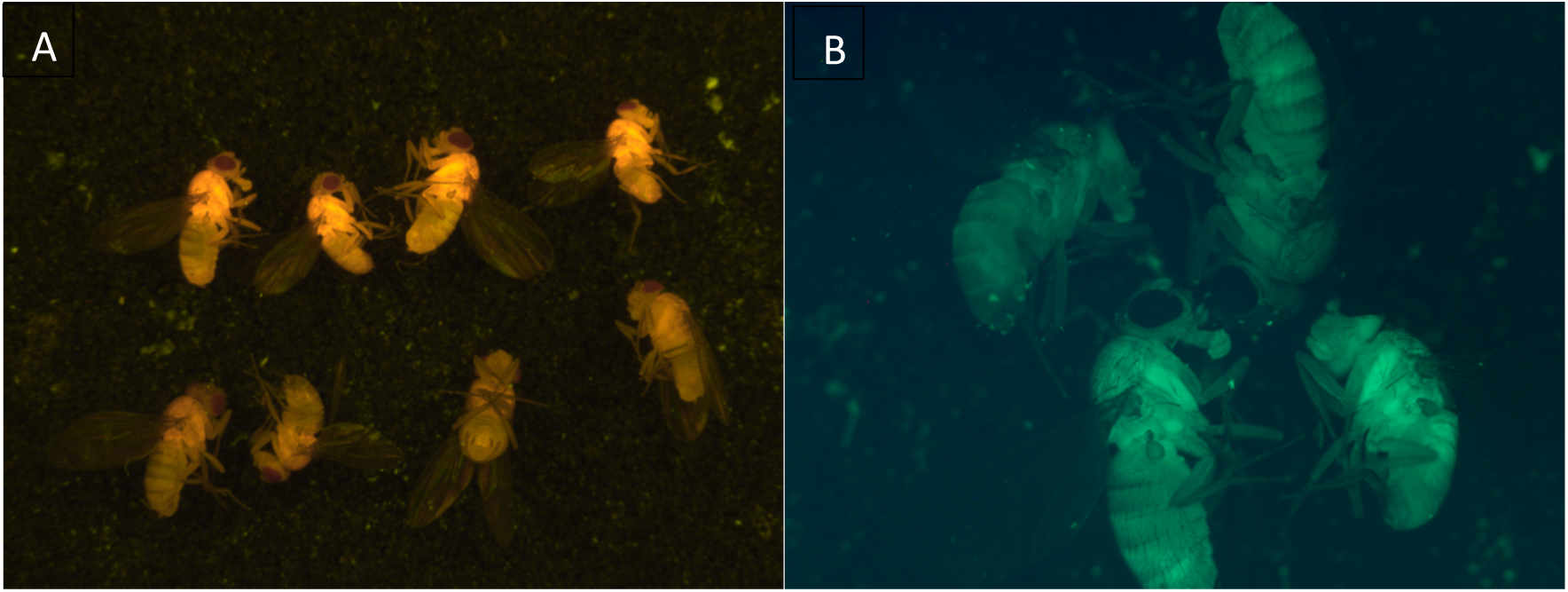
Heterozygous vs. Homozygous transgenic *Drosophila* carrying UAS-Gal4 and UAS-Gal80 transgenes. (A) Flies in the top row are homozygous for the transgenes while flies in the bottom row are heterozygous. The homozygotes can be identified and separated by their brighter fluorescence intensity compared to heterozygotes. (B) *D. melanogaster* under the narrow band green filter. The top two flies are heterozygous and the bottom two are homozygous as identified by their brighter fluorescence intensity.

**Fig S4.**
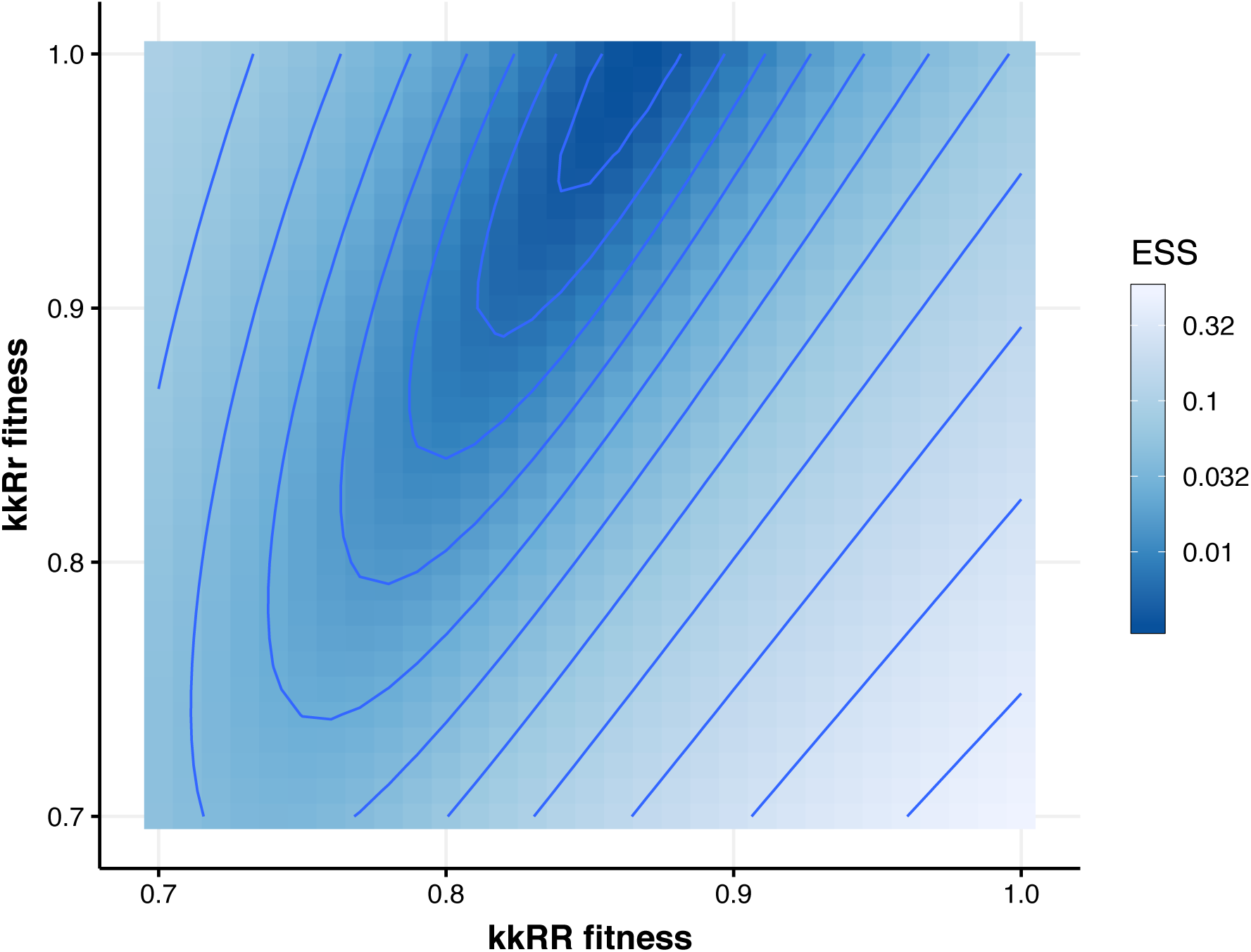
Fitness parameter estimation for the Gal80 Rescue experiment. Color and contour lines show the log error sum of squares (ESS) for each combination of kkRR and kkRr fitnesses. A fitness of 1 is equivalent to wildtype. The model simulation is most consistent with the data when kkRr=1 and kkRR=0.87 (dark blue). There is a region of parameter space for which the goodness of fit is roughly similar, with a slight decrease in both kkRr and kkRR fitnesses maintaining a good fit.

**Table S1.** Fitness parameters used in model simulations for each genotype.

| Genotype | DSCP-Gal4;Gal80 | Hsp70-Gal4;Gal80 | Hsp70-Gal4-hid; Gal80 | Gal80 |
| --- | --- | --- | --- | --- |
| kkrr (wt) | 1 | 1 | 1 | 1 |
| kkRR | 0.87 | 1 | 1 | 0.87 |
| kkRr | 1 | 0.78 | 0.68 | 1 |
| KkRR | 0.83 | 0.67 | 0.86 | - |
| KkRr | 0.77 | 0.18 | 0.15 | - |
| KKRR | 0.70 | 0 | 0 | - |
| KKRr | 0.73 | 0 | 0 | - |
| KKrr | 0 | 0 | 0 | - |
| Kkrr | 0 | 0 | 0 | - |

**Table S2.** Drosophila lines used in this research. The name of line, genotype, source (Bloomington Drosophila Stock Center or publication (1,2,3,4)) and use in this research are listed.

| Line | Genotype | Source | Use in this research |
| --- | --- | --- | --- |
| attP40 | y<sup>1</sup>v<sup>1</sup>P{y[+t7.7]=nos-phiC31\int.NLS}X;<br>P{y[+t7.7]=CaryP}attP40 (II) | BDSC # 25709 | Location for Rescue Insertion |
| attP2 | y<sup>1</sup>sc<sup>1</sup>v<sup>1</sup>P{y[+t7.7]=nos-phiC31\int.NLS}X;<br>P{y[+t7.7]=CaryP}attP2 (III) | BDSC # 25710 | Location for Killer Insertion |
| attP9A | y<sup>1</sup>w<sup>*</sup>P{y[+t7.7]=nos-phiC31\int.NLS}X;<br>PBac{y[+]-attP-9A}VK00027 (III) | BDSC # 35569 | Location for Killer Insertion |
| UAS-RedStinger | P{w[ +mC]=UAS-RedStinger}4 | (1) | Gal80 Repression Experiment |
| GMR-Gal4 | w[1118];P{y[+t7.7]w[+mC]=GMR-Gal4}attP2 | (2) | Gal80 Repression Experiment |
| Actin-Gal4 | w[1118];P{y[+t7.7]w[+mC]=Act-Gal4}attP2 | (3) | Gal80 Repression Experiment |
| Elav-Gal4 | w[1118];P{y[+t7.7]w[+mC]=Elav-Gal4}attP2 | (4) | Gal80 Repression Experiment |

**Table S3.**
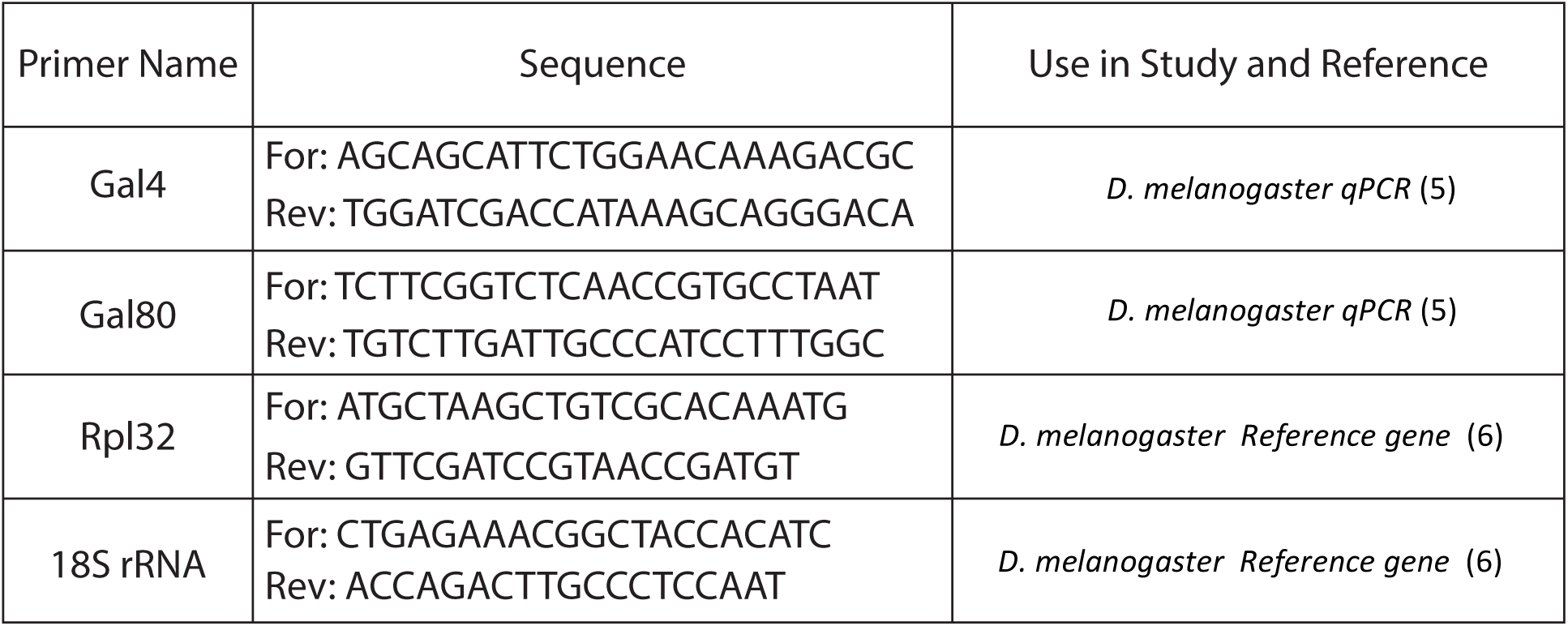
List of primer sequences, target and reference genes, used in qRT-PCR experiments (5,6).

## Acknowledgements

We thank Fred Gould and Alun Lloyd for helpful discussions. We thank Rob Harrell, Channa Aluvihare, Azadeh Aryan, Zach Adelman and Esther Belikoff for training in insect microinjections. SHW was supported by NSF IGERT Program grant-1068676. MRV received support from the Research Training Group in Mathematical Biology, funded by a National Science Foundation grant RTG/DMS-1246991.

